# Activation of G proteins by guanine nucleotide exchange factors relies on GTPase activity

**DOI:** 10.1101/032607

**Authors:** Rob J Stanley, Geraint MH Thomas

## Abstract

G proteins are an important family of signalling molecules controlled by guanine nucleotide exchange and GTPase activity in what is commonly called an ‘activation/inactivation cycle’. The molecular mechanism by which guanine nucleotide exchange factors (GEFs) catalyse the activation of monomeric G proteins is well-established, however the complete reversibility of this mechanism is often overlooked. Here, we use a theoretical approach to prove that GEFs are unable to positively control G protein systems at steady-state in the absence of GTPase activity. Instead, positive regulation of G proteins must be seen as a product of the competition between guanine nucleotide exchange and GTPase activity – emphasising a central role for GTPase activity beyond merely signal termination. We conclude that a more accurate description of the regulation of G proteins via these processes is as a ‘balance/imbalance’ mechanism. This result has implications for the understanding of many intracellular signalling processes, and for experimental strategies that rely on modulating G protein systems.

## Introduction

G proteins are an important and universal family of intracellular signalling molecules, incorporating both the alpha subunits of heterotrimeric G proteins and the Ras small monomeric G proteins. Most G proteins bind guanine nucleotides (GDP, GTP) in a strongly conserved nucleotide binding pocket – an ancient mechanism preserved in both eukaryotes and prokaryotes (Simon et al. 1991; Dong et al. 2007; Rojas et al. 2012). Typically, G proteins transition between two discrete conformations with distinct signalling functions depending on which nucleotide is bound, and so G proteins are often referred to as ‘molecular switches’. G protein regulatory systems are crucial components of many intracellular processes – incorrect regulation of G proteins has been implicated in disease: cancer (Young et al. 2009; Vigil et al. 2010; O’Hayre et al. 2013), cardiovascular disease (Loirand et al. 2013), genetic disorders (Seixas et al. 2013), among many others.

Regulation of G protein activation is largely controlled by two mechanisms (Figure 1A) and is commonly described as an ‘activation/inactivation cycle’ between the GTP-bound ‘on/active’ state and the GDP-bound ‘off/inactive’ state (Vetter and Wittinghofer 2001; Oldham and Hamm 2008). Activation of G proteins is controlled by accessory proteins which catalyse guanine nucleotide exchange – the sequential release of GDP and binding of GTP. For monomeric G proteins these are known as guanine nucleotide exchange factors (GEFs). For heterotrimeric G proteins, G protein coupled receptors (GPCRs) fulfil this role. Inactivation of G proteins is controlled by GTPase activity which may either be intrinsic, or be provided via accessory GTPase-activating proteins (GAPs). It is generally thought that GTPase activity is required for the termination of G protein signalling but that it is not essential for signal transmission (Takai et al. 2001).

**Figure 1.**
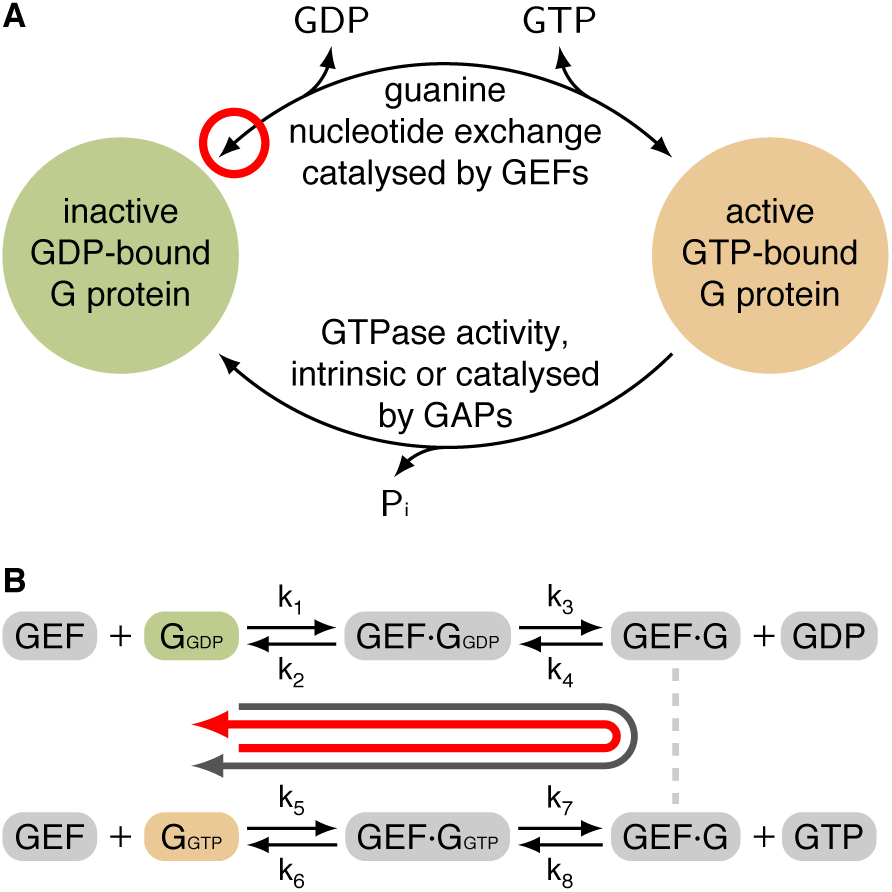
The activation of G proteins is regulated by GEFs and GTPase activity. **A** G proteins are controlled by GEFs which catalyse the sequential release and binding of guanine nucleotides, and by GTPase activity (both intrinsic and GAP-mediated) which hydrolyses the bound GTP to form GDP. The red circle highlights that the GEF mechanism is completely reversible. **B** The reversible mechanism by which a GEF catalyses guanine nucleotide exchange on a G protein proceeds through a series of GEF·G protein complexes (Bos et al. 2007). Parameters *k_i_* are kinetic rates which are unique to each G protein:GEF system. Associated species (free GEF, GTP, GDP) have not been drawn. The grey arrow identifies forwards nucleotide exchange, catalysing the activation of the G protein. The ref arrow identifies reverse nucleotide exchange, catalysing the inactivation of the G protein.

An often overlooked property of GEFs is that their catalytic mechanism is completely reversible (Figure 1B) (Goody 2014). GEF-binding is not specific to GDP-bound G protein – GEFs can also bind to GTP-bound G protein and catalyse the reverse nucleotide exchange, GTP to GDP. In this way, GEFs are capable of inactivating G proteins (Bos et al. 2007). The extent to which the reversibility of this mechanism has been overlooked is demonstrated by the sheer number of publication which include diagrams where arrows corresponding to GEF-mediated regulation are drawn as unidirectional – missing the reverse arrowhead highlighted in Figure 1A. This error is perhaps best illustrated by its occurrence in core biology textbooks, for example:

- Figures 3–66 and 3–68 in Alberts et al. (2014)
- Figures 16–15 and 16–16 in Alberts et al. (2013)
- Figure 4, box 12–2 in Nelson and Cox (2013)
- Figure 13.40 in Berg et al. (2010)
- Figure 19–40 in Voet and Voet (2010)
- Figure 7.12A in Hancock (2010)
- Figure 10.3 and 10.4 in Bolsover et al. (2011)
- Figure 42.4 in Baynes and Dominiczak (2014)

There has been recent renewed interest in understanding the roles and functions of GEFs based on a proper consideration of their enzyme kinetics (Northup et al. 2012; Randazzo et al. 2013; Goody 2014). Here we develop the theoretical understanding of G protein regulation by GEFs and GTPase activity through exploring the consequences of the reversibility of the GEF mechanism. We use mathematical methods to investigate G protein regulatory systems independent of measured kinetic rates, in the context of the physiologically important steady-state dynamics. This allows us to comment and draw conclusions on the qualitative behaviours of G protein:GEF:GTPase systems under a wide variety of conditions.

## Results

### Qualitative differences between reversible and irreversible mechanisms

To demonstrate the qualitative difference between a reversible and an irreversible mechanism we derived mass-action models of the GEF mechanism (Figure 1B, Methods) and an artificial irreversible mechanism generated by disallowing release of GTP from the G protein·GEF complex.

The reversible and irreversible models were simulated: in the absence of GTPase activity (Figures 2A, 2D); with intrinsic GTPase activity, modelled by exponential decay (Figures 2B, 2E); and with GAP-mediated GTPase activity, modelled using the Michaelis-Menten equation (Figures 2C, 2F). To ensure that simulations were physiologically plausible, kinetic rates measured for the the Ran:RCC1 system were used (Klebe et al. 1995). A GTP:GDP ratio of 10:1 was used to emulate the relative levels in eukaryotic cells.

**Figure 2.**
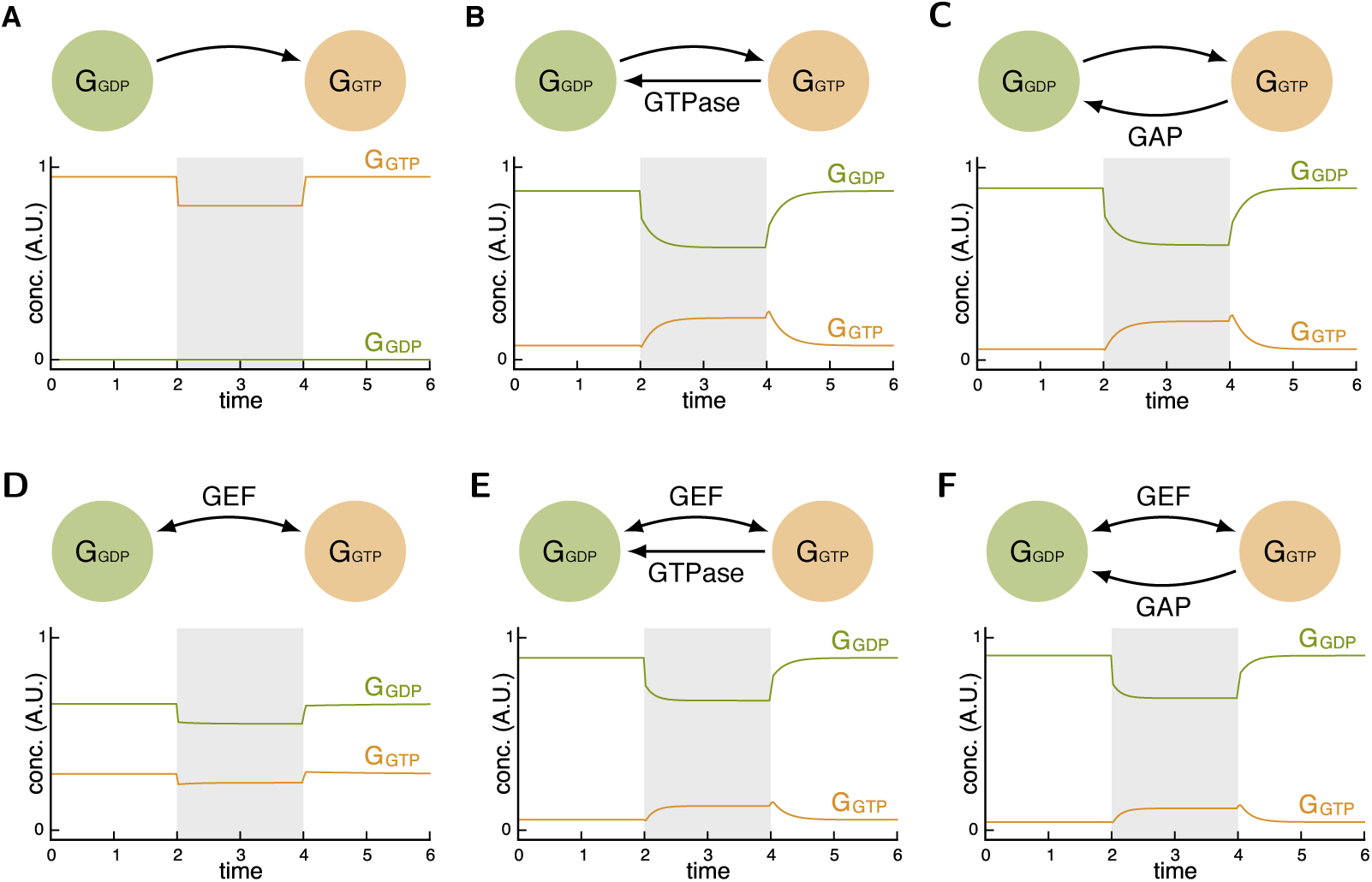
Apparent activation of G proteins via GEFs is only observed when GTPase activity is present. Simulation of mass-action models, using parameters described in Table S1, and where *G*_GXP_ denotes GXP-bound G protein. Where indicated as present, intrinsic GTPase activity was modelled as exponential decay, GAP-mediated GTPase activity by the Michaelis-Menten equation. The shaded region denotes stimulation of the system through increasing the active GEF 4-fold from its basal concentration. For all simulations, steady-state concentrations were used as the initial conditions. Mass corresponding to GEF·G protein complexes has not been drawn. **A, B, C** An artificial irreversible model, constructed by assuming the rate of release of GTP from the active G protein·GEF complex is zero. **D, E, F** The reversible GEF mechanism (Figure 1B).

In the presence of either form of GTPase activity both reversible and irreversible mechanisms display similar behaviour which is consistent with observations of GEF-mediated activation of G proteins in a wide range of biological systems (Janetopoulos et al. 2001; Peyker et al. 2005; Adjobo-Hermans et al. 2011; Chang and Ross 2012; Oliveira and Yasuda 2013).

In the absence of GTPase activity we see a qualitative difference in the behaviour of the two mechanisms; each distinct from their shared behaviour in the presence of GTPase activity. While both mechanisms show an inhibitory effect (which will discussed below in more detail for the GEF mechanism), the steady-state concentrations of active and inactive G protein differ substantially. Through this example we demonstrate how the assumption of an irreversible model would lead to incorrect conclusions when considering extremal (i.e. diseased) states.

### GEFs act to attain a constant ratio of inactive to active G protein

We derived a simplified quasi-steady-state model of the GEF mechanism (Figure 1B) in an equivalent manner to the derivation of the Michaelis-Menten equation (Michaelis and Menten 1913; Briggs and Haldane 1925; Johnson and Goody 2011; Gunawardena 2012). This quasi-steady-state model captures the behaviour of a generic G protein regulatory system in a single equation:

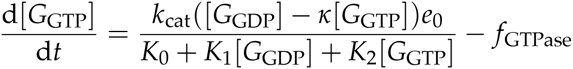

Here [*G*_GXP_] is the concentration of GXP-bound G protein and *κ* is the ratio of the backwards to the forwards kinetic rates. (For definitions of the other parameters see the Methods section.)

At steady-state (setting the above equation equal to zero), in the absence of GTPase activity, we find that the ratio of inactive to active G protein must always equal the value of the constant *κ*. An equivalent statement is: GEFs act to produce a constant proportion of active G protein. While the ratio of inactive to active G protein (*κ*) and proportion of active G protein (1/*_k_* _+ 1_) will vary for different G protein:GEF systems, these values will remain constant within a system, independent of the G protein or GEF concentrations.

### GEFs can be inhibitory

The commonly used description of GEFs as ‘activators’ of G proteins is contradicted by the inhibitory effect seen when the GEF mechanism is simulated in the absence of GTPase activity (Figure 2D). This demonstrates the inadequacy of this description.

The inhibitory effect can be explained by an equivalent increase in the concentrations of intermediate G protein·GEF complexes. Values for the concentrations of these intermediate complexes were derived as part of the construction of the quasi-steady-state model. Using these values, we obtained an equation for the proportion of (free) active G protein in terms of the total concentration of GEF. This equation is plotted with the rates described for the Ran:RCC1 system in Figure 3A. Using this equation we are able to prove that in the absence of GTPase activity the concentration of active G protein is inversely related to the total concentration of GEF. As the concentration of GEF increases, the concentration of G protein will always decreases, and vice-versa.

**Figure 3.**
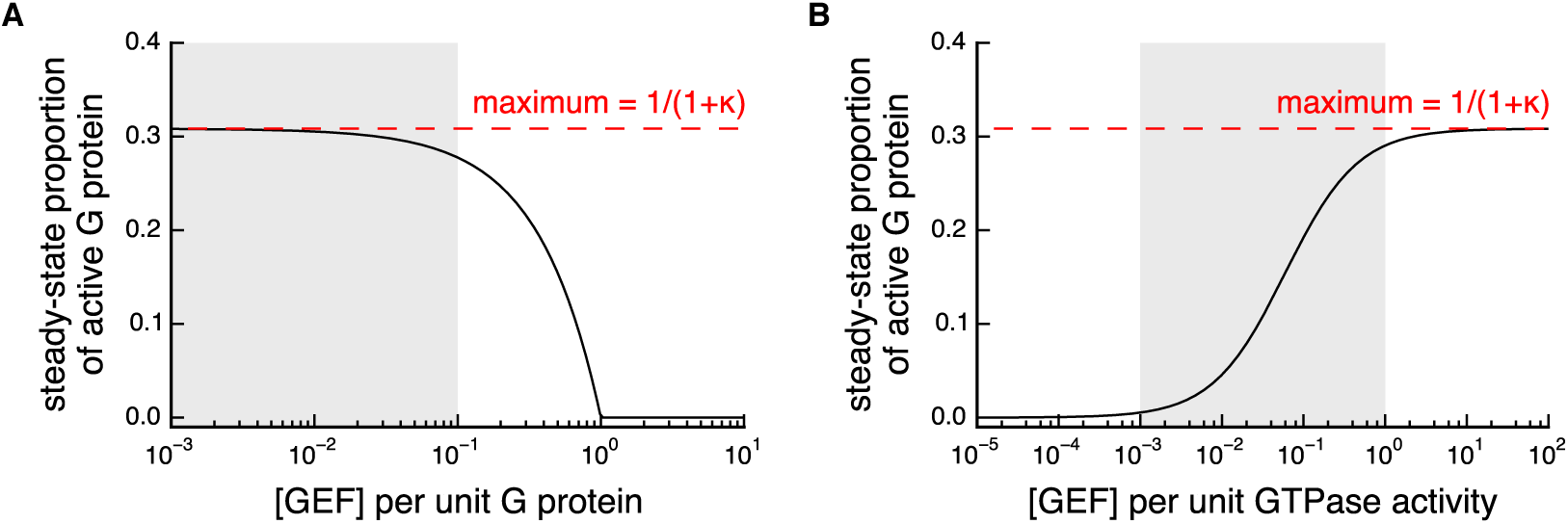
GTPase activity restores the ability of GEFs to positively regulate a G protein by moving the system away from equilibrium. The relationship between the concentration of GEF and the steady-state proportion of active G protein (equation (5), equation (7)) illustrated using parameters described for the Ran:RCC1 system (Klebe et al. 1995) and unit concentration of G protein. The activation cannot be increased above a theoretical maximum equilibrium value derived from the ratio of the total forwards and backwards catalytic rates of the GEF (*κ*). The shaded region denotes the region which is most likely to be physiologically relevant. **A** In the absence of GTPase activity (equation (5)), increasing the GEF concentration can only decrease the steady-state concentration of active G protein, instead producing irrelevant GEF·G protein complexes. **B** In the presence of GTPase activity (equation (7)), the steady-state concentration of active G protein is suppressed. Increasing the (relative) concentration of GEF acts to counter this suppression, driving the activation state back towards the maximum equilibrium value.

Note that a high concentration of GEF will also lead to a faster total catalytic rate (a larger *V*_max_). This suggests that there will be a tradeoff in terms of increasing the concentration of GEF: a low concentration of GEF means that there will be little inhibition, but a slow total rate; a high concentration of GEF will lead to inhibition, but a fast total rate. We therefore hypothesise that for a healthy G protein system, the concentration of GEF will lie in a physiologically relevant region, where the inhibitory effect is not so pronounced, but where there is still sufficient GEF to catalyse nucleotide exchange at an appropriate rate.

### GTPase activity has a functional role in the observed activation of G proteins

The simulations of the GEF mechanism show that GTPase activity is sufficient to restore an apparent GEF-mediated activation (Figures 2E, 2F). By comparing these with the simulation of the system without GTPase activity (Figure 2D), we can see how this activation arises. Initially, due to the GTPase activity, the activation state reached by the system is suppressed – it is much reduced from the activation state reached in the absence of GTPase activity. An increase in the concentration of GEF is then able to positively regulate the system by moving the activation state closer to the activation state reached in the absence of GTPase activity (even though this state may itself be reduced).

For intrinsic GTPase activity we obtained an equation which describes the effect of the relative rates of GEF-catalysed nucleotide exchange and GTPase activity on the proportion of G protein which is active. This equation is plotted with example parameters in Figure 3B, where we see a sigmoidal response such that increasing the concentration of GEF (relative to the GTPase activity) increases the concentration of active G protein. Again this allows us to hypothesise that, for a healthy G protein system, the relative rates of nucleotide exchange and GTPase activity must lie in this sigmoidal region, in order for the system to properly respond to an activating or inhibitory signal.

Together, this clearly demonstrates a requirement for GTPase activity for the observable activation of G proteins by GEFs. The proposed mechanism of regulation for a generic G protein:GEF:GTPase system can be summarised as follows: 1. GTPase activity inactivates the G protein system by altering the ratio of inactive to active G protein away from a GEF-mediated equilibrium. 2. If the rate of guanine nucleotide exchange increases or the GTPase activity decreases, the proportion of active G protein will then move towards the GEF-mediated equilibrium, generating an observed activation.

## Discussion

We have shown that there are certain universal properties of GEF-mediated regulation of G proteins that arise from the reversibility of its mechanism and which are independent of specific kinetic rates. The complete reversibility of the GEF mechanism means that at steady-state any GEF acts to produce a constant ratio of inactive to active G protein – giving a theoretical maximum proportion of active G protein. Once this maximum is attained, then any subsequent increase in the concentration of GEF—the ‘activator’ of the system—cannot increase the concentration of active G protein. Instead this will lead to inhibition caused by creation of excess intermediate G protein·GEF complexes.

We urge caution against naïve description of GEFs as ‘enzymes that activate G proteins’ and against representations that show this mechanism as irreversible as we have shown how these shorthands distort our understanding of the underlying biology. We have demonstrated that GEFs should not be described as enzymes that convert a substrate into product, but as enzymes that act to attain an equilibrium—a balance—of active and inactive G protein. The two key roles of GTPase activity are then: to drive the system away from this equilibrium—to create an imbalance—and so permit positive regulation by GEFs; and to confer a unique directionality on the G protein regulatory ‘cycle’. Therefore we suggest that G protein signalling controlled by GEFs and GTPase activity should not be described as an ‘activation/inactivation’ cycle but rather as a system that is controlled through ‘regulated balance/imbalance’.

Both the complete reversibility of guanine nucleotide exchange and associated requirement for GTPase activity as a functional component in the activation of G proteins has previously been under-appreciated. This may be due to the almost exclusive use of experimental systems where the GDP form of the G protein is the unique starting condition and where uptake of GTP is monitored as the GEF assay. We also note that our simulations show that an artificial irreversible mechanism (Figures 2B, C) and reversible GEF mechanism (Figures 2E, F) have similar profiles in the presence of GTPase activity and so under many conditions it may be difficult to experimentally distinguish these mechanisms.

We predict that experimental protocols which attempt to regulate G proteins by the over-expression of a GEF are likely to produce unexpected behaviour. We expect that in many cases this may cause inhibition of the G protein rather than activation (Figure 3A). Activation of G proteins should therefore be preferentially targeted by reduction of the relevant GTPase activity (Figure 3B). Note that these results remain consistent with the long-established use of dominant negative mutants for the inhibition of G protein systems (Feig 1999; Barren and Artemyev 2007). We accept that many previous studies that have ignored the reversibility of GEFs will have made conclusions that are valid under many conditions. But we stress that in extremal scenarios (such as in disease) those conclusions may not always hold.

Additionally, we hope that this new perspective in considering the control of G proteins will lead to novel approaches for the control of G protein systems. GEFs have previously been suggested as potential therapeutic targets (Bos et al. 2007). Our results extend this to a novel, and seemingly paradoxical, mechanism by which over-expression of an activator could lead to the inhibition of its substrate. This may have implications in G protein systems with diminished GTPase activity, for example constitutively active transforming mutations in Ras common in cancers (Stephen et al. 2014), where additional GAP activity would have no effect but where sequestration of active G protein by a GEF may be useful alternative.

The mathematical underpinning to our results mean that they should hold for any G proteins:GEF system so long as the mechanism is consistent with that studied here (Figure 1A), and under the reasonable assumption that the majority of its functional signalling is due to the steady-state behaviour. The precise tradeoffs for any system (equilibrium ratios, total rates, and scale of inhibition) will depend on the specific kinetic rates for the GEF and the strength of GTPase activity, but the overall qualitative characteristics should remain consistent across all such systems. Conclusions based on alternative mechanisms, for instance systems with an implicit G protein·GEF·GAP complex (Berstein et al. 1992), would require further analysis.

## Methods

### The following mathematical analysis uses the notation

- G protein without nucleotide bound → *G*
- G protein with GDP bound → *G*_GDP_
- G protein with GTP bound → *G*_GTP_
- GEF → *E*

The volume concentration of a species *S* will be denoted by [*S*].

### Mass-action model

A deterministic ordinary differential equation (ODE) model of the GEF mechanism (Figure 1B) was derived using the law of mass-action:

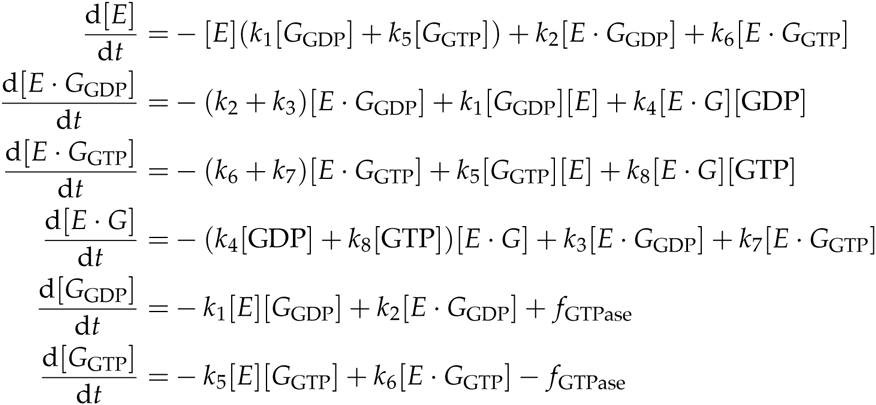

We assume: for systems with no GTPase activity, *f*_GTPase_ = 0; for systems with intrinsic GTPase activity, *f*_GTPase_ = *k*_ase_[*G*_GTP_]; and for systems with GAP-mediated GTPase activity, 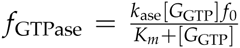where *f*_0_ is the total concentration of GAP.

There is an equation for the conservation of mass of GEF:

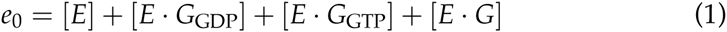

And an equation for the conservation of mass of G protein:

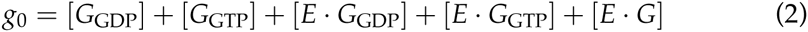

### Simulation of the mass-action model

The parameters used for the simulations in Figure 2 are summarised in Table S1. Wherever possible, parameters measured for the Ran:RCC1 system were used (Klebe et al. 1995). The irreversible model was generated by setting *k*_7_ = 0. (Alternative irreversible models could be generated by setting any one or more of the reverse reaction rates to zero.)

All simulations were started from steady-state and generated by numerical integration of the mass-action equations, with the exception of free enzyme concentration [*E*] which was calculated from the total mass of enzyme equation (1) with:

- *e*_0_ = 0.05 during 0 ≤ *t <* 2
- *e*_0_ = 0.2 during 2 ≤ *t <* 4
- and free GEF (*E*) removed from the simulation until *e*_0_ = 0.05 during *t* ≥ 4

### Quasi-steady-state model

Quasi-steady-state solutions for the intermediate enzyme complexes of the GEF mechanism (Figure 1B) were derived using the framework of Gunawardena (2012) (Figure S1):

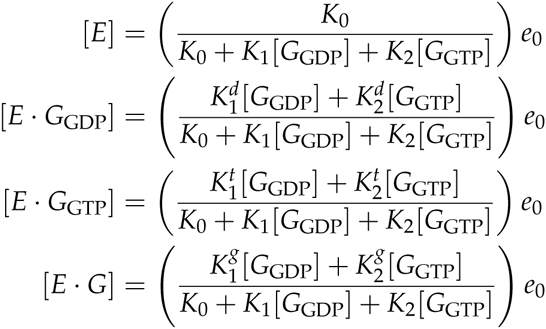
where the 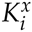and the *K_i_* are summary parameters (defined in Table S1).

These quasi-steady-state solutions were substituted into the equation for the rate of change of [*G*_GTP_] given in the mass-action model, to obtain a quasi-steady-state model for a generic GEF acting on a generic G protein:

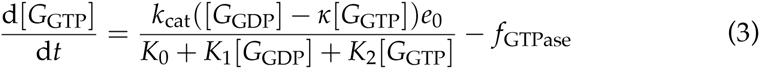

where *k*_cat_ is the forward catalytic rate; *κ* is the ratio of the backwards to the forwards kinetic rates, multiplied by the ratio of GDP to GTP.

This equation does not consider mass held in G protein·GEF intermediate complexes and so is only a good approximation when *e*_0_ *≪ g*_0_. Note that with *f*_GTPase_ = 0 this model reduces to the Michaelis-Menten equation when *y* = 0, and is equivalent to the equation used by Randazzo et al. (2013) when the concentration of GTP is absorbed into the summary paramters.

### Steady-state ratio of inactive to active G protein

At steady-state with *f*_GTPase_ = 0, equation (3) implies:

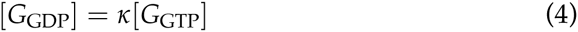

Assuming that *e*_0_ *≪ g*_0_, equation (2) simplifies to *g*_0_ = [*G*_GDP_] + [*G*_GTP_], into which equation (4) can be substituted to obtain:

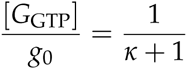

This is the maximum steady-state proportion of active G protein.

### Active G protein as a function of GEF concentration (without GTPase activity)

The effect of increasing the concentration of GEF on the steady-state concentration of active G protein in the absence of GTPase activity ( *f*_GTPase_ = 0) was investigated.

The quasi-steady-state solutions for the intermediate enzyme complexes and equation (4) were substituted into equation (2) to obtain:

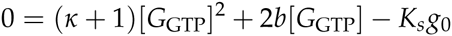
where 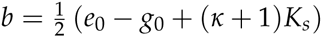and 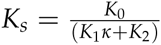

This quadratic equation has one positive solution:

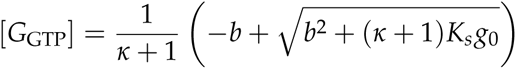

Alternatively, the proportion of active G protein is:

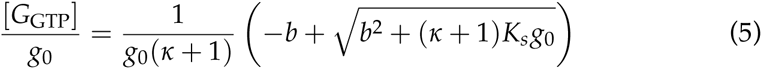

We are interested in the rate of change of [*G*_GTP_] with respect to *e*_0_, the total concentration of GEF. As *b* (and only *b*) is a function of *e*_0_, we can examine:

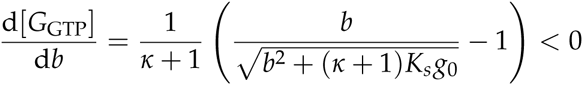

As this equation is always negative, the concentration of active G protein must decrease as the concentration of GEF is increased (and vice-versa).

### Active G protein as a function of GEF concentration (with GTPase activity)

The effect of increasing the concentration of GEF on the steady-state concentration of active G protein with GTPase activity ( *f*_GTPase_ = *k*_ase_[*G*_GTP_]) was investi gated.

At steady-state 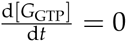implies:

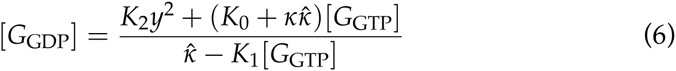

where 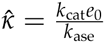

Again assuming that *e*_0_ *≪ g*_0_, equation (2) simplifies to *g*_0_ = [*G*_GDP_] + [*G*_GTP_], into which equation (6) can be substituted to obtain:

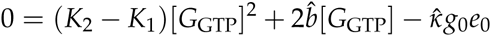
where 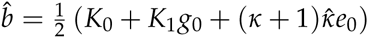.

This quadratic equation has one solution that lies in the region 0 ≤ [*G*_GTP_] ≤ *g*_0_:

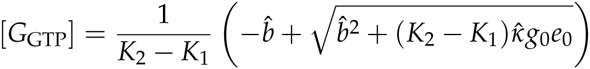

Alternatively, the proportion of active G protein is:

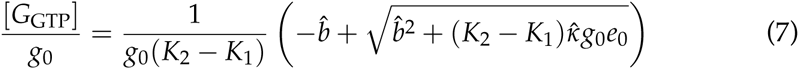

This equation describes the steady-state concentration of active G protein as a function of 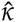, the ratio of the rate of forwards GEF-mediate nucleotide exchange to the rate of GTPase activity.

## Acknowledgements

We would like to thank the members of the UCL CSSB lab including Janine Symonds and Chris Barnes, and also Tina Daviter for their critical reading of previous versions of the manuscript. We would like to thank Kevin Bryson for his support. Funding for this study was provided by the British Heart Foundation.

## Author contributions

This work formed part of the doctoral research of RS under the supervision of GT. RS produced mathematical results, and GT and RS wrote the paper.

---

**Figure S1.**
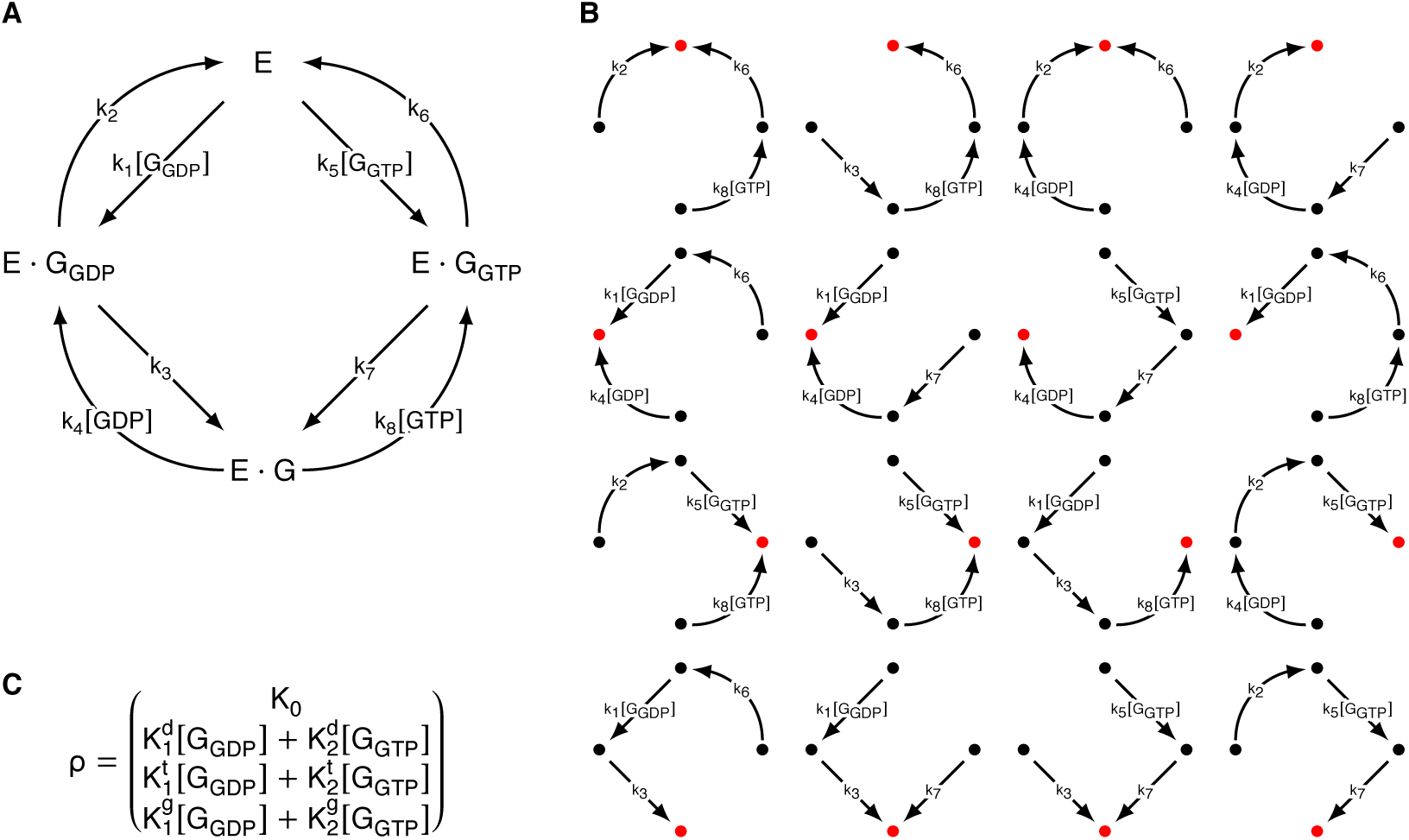
Application of the framework of Gunawardena (2012) to the mechanism for the GEF mediated release and binding of guanine nucleotides to G proteins. **A** The graph on the enzyme complexes with complexes as vertices and edges representing reactions labelled by rates and partner species. **B** All possible directed spanning trees of the graph on the enzyme complexes. The red vertex denotes the root of each spanning tree. **C** The basis element, *ρ*, generated from the each spanning trees: the sum over each root vertex, of the products of the labels of each spanning tree. Every steady-state of the original system *X* = ([*E*], [*E·G*_GDP_], [*E·G*_GTP_], [*E·G*])*^T^* is a solution to the equation *X* = *λρ* where *λ* is a constant. We manipulate this equation to obtain 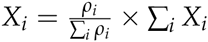

**Table S1.** Concentrations, kinetic parameters, and summary parameters used for Figures 2 and 3. Where applicable, the definitions of the summary parameters in terms of the individual kinetic parameters are stated. Value obtained from (Klebe et al. 1995).

| Rate | Reversible model | Irreversible model | Definition |
| --- | --- | --- | --- |
| $f_0$ | 1 | 1 | |
| [GTP] | 10 | 10 |  |
| [GDP] | 1 | 1 |  |
| $k_1^*$ | $7.4 \times 10^7$ | $7.4 \times 10^7$ | |
| $k_2^*$ | 55 | 55 | |
| $k_3^*$ | 21 | 21 | |
| $k_4^*$ | $1.1 \times 10^7$ | $1.1 \times 10^7$ | |
| $k_5^*$ | $1.0 \times 10^8$ | $1.0 \times 10^8$ | |
| $k_6^*$ | 55 | 55 | |
| $k_7^*$ | 19 | 0 | |
| $k_8^*$ | $0.6 \times 10^6$ | $0.6 \times 10^6$ | |
| k_ase | 4 | 4 |  |
| K_m | 0.7 | 0.7 |  |
| K_ic | 100 | 100 |  |
| $K_1^d$ | $8.466 \times 10^{16}$ | $6.919 \times 10^{16}$ | $k_1(k_6k_8[\text{GTP}] + k_4(k_6 + k_7)[\text{GDP}])$ |
| $K_2^d$ | $2.090 \times 10^{16}$ | 0 | $k_4k_5k_7[\text{GDP}]$ |
| $K_1^g$ | $1.150 \times 10^{11}$ | $8.547 \times 10^{10}$ | $k_1k_3(k_6 + k_7)$ |
| $K_2^g$ | $1.444 \times 10^{11}$ | 0 | $k_5k_7(k_2 + k_3)$ |
| $K_1^t$ | $9.324 \times 10^{15}$ | $9.324 \times 10^{15}$ | $k_1k_3k_8[\text{GTP}]$ |
| $K_2^t$ | $1.061 \times 10^{17}$ | $1.061 \times 10^{17}$ | $k_5(k_8(k_2 + k_3)[\text{GTP}] + k_2k_4[\text{GDP}])$ |
| $K_0$ | $6.985 \times 10^{10}$ | $5.836 \times 10^{10}$ | $k_6k_8(k_2 + k_3)[\text{GTP}] + k_2k_4(k_6 + k_7)[\text{GDP}]$ |
| $K_1$ | $9.398 \times 10^{16}$ | $7.851 \times 10^{16}$ | $K_1^d + K_1^g + K_1^t$ |
| $K_2$ | $1.270 \times 10^{17}$ | $1.061 \times 10^{17}$ | $K_2^d + K_2^g + K_2^t$ |
| $k_{\text{cat}}$ | $5.128 \times 10^{17}$ | $5.128 \times 10^{17}$ | $k_1k_3k_6k_8[\text{GTP}]$ |
| $\kappa$ | 2.242 | 0 | $\frac{k_2k_4k_5k_7[\text{GDP}]}{k_1k_3k_6k_8[\text{GTP}]}$ |
| $K_s$ | $2.069 \times 10^7$ | $5.500 \times 10^{-7}$ | $\frac{K_0}{(K_1\kappa + K_2)}$ |

